# Mass recalibration for desorption electrospray ionization mass spectrometry imaging using endogenous reference ions

**DOI:** 10.1101/2021.03.29.437482

**Authors:** Paolo Inglese, Helen Xuexia Huang, Vincen Wu, Matthew R Lewis, Zoltan Takats

## Abstract

1

**Background:** Mass spectrometry imaging (MSI) data often consist of tens of thousands of mass spectra collected from a sample surface. During the time necessary to perform a single acquisition, it is likely that uncontrollable factors alter the validity of the initial mass calibration of the instrument, resulting in mass errors of magnitude significantly larger than their expected values. This phenomenon has a two-fold detrimental effect: a) it reduces the ability to interpret the results based on the observed signals, b) it can affect the quality of the observed signal spatial distributions.

**Results:** We present a post-acquisition computational method capable of reducing the observed mass drift by up to 60 ppm in biological samples, exploiting the presence of typical molecules with a known mass-to-charge ratio. The procedure, tested on time-of-flight (TOF) and Orbitrap mass spectrometry analyzers interfaced to a desorption electrospray ionization (DESI) source, improves the molecular annotation quality and the spatial distributions of the detected ions.

**Conclusion:** The presented method represents a robust and accurate tool for performing post-acquisition mass recalibration of DESI-MSI datasets and can help to increase the reliability of the molecular assignment and the data quality.

## 2 Introduction

In the last twenty years, mass spectrometry imaging (MSI) has attracted increasing attention as a technology capable of capturing molecular spatial patterns from sample surfaces. Among the available ionization sources, such as matrix-assisted laser desorption^1^ (MALDI) or secondary ion mass spectrometry^2^ (SIMS), desorption electrospray ionization^3^ (DESI) has gained popularity and been widely applied thanks to its relatively simpler preparation process. A map of the detected ions’ spatial distribution is usually represented as an image and analyzed for identifying informative spatial patterns. MSI has been successfully deployed in various application areas, from cancer research^4–7^ to pharmacology.^8–12^ However, as an emerging technology, MSI still faces implementation challenges for controlling the quality of data produced.^13–15^ The accuracy of mass-to-charge ratio (*m/z)* measurements under sustained use are chief among these concerns.

MSI acquisitions often consist of tens of thousands of spectra (pixels), corresponding to several hours of operation, during which mass measurements can be subject to substantial drift. For example, a small image of 10 mm x 10 mm, acquired at a running speed of 1 scan/s, with a spatial resolution of 100 µm, requires about 2.8 hours. Shifting mass measurements are detrimental to the analyzed spatial patterns and can result in erroneous chemical annotations. Additionally, as MSI relies on only a single dimension of separation for detected ions, errors of a few parts per million (ppm) can result in hundreds of candidate molecular identities, interpreting in terms of biochemical hypotheses a practical challenge.

For these reasons, it is crucial to develop quality control approaches for MSI that mitigate this issue and yield accurate *m/z* measurements with high precision during extended instrument use. Solutions successfully applied in more routine hyphenated techniques (e.g., liquid chromatography-mass spectrometry LC-MS) include periodic recalibration or more frequent calibration based on the measurement of one or more standard reference materials during the acquisition. Such an approach is infeasible for the current MSI technology since the acquisition is performed continuously while the probe moves across the sample surface. However, this approach may be emulated by adding the standard material to the sample. ^16^ Although this approach can be used to facilitate post-acquisition *m/z* correction of mass accuracy, concerns may arise about the unknown effect on the ionization efficiency caused by these exogenous molecules’ presence. In the specific case of DESI-MSI, usually, only one reference molecule (lock mass) is added to the solvent, which can be insufficient to capture the non-linear drift across the measured *m/z* range.

Another class of approaches relies on using endogenous or background peaks as candidate reference ions to estimate the mass shift occurring in each spectrum and map the measured *m/z* onto their corresponding recalibrated values.^17^ This class of methods requires prior knowledge about the molecular content of the sample.

Boskamp et al. showed that endogenous signal (chemical noise) could improve the mass accuracy in MALDI-MSI of peptide datasets.^18^ Recently, La Rocca et al. have presented an algorithm to recalibrate MSI datasets using a linear fit on the observed mass errors from endogenous biological peaks.^19^ In their method, they treated each spectrum as independent.

Here, we present a different approach based on a fixed set of reference ions across the entire MSI dataset, in which the individual mass shifts are modeled as a time series. The advantages of global references are that they 1) define a fixed boundary for the *m/z* extrapolation, 2) improve evaluation of the quality of the matching procedure by visualizing their spatial distribution, and 3) increase the robustness of the matching process as a time-dependent model.

The approach leverages the ubiquity of common structural components of tissues as dependable sources of reference values, using them as multiple points to estimate error and correct all observed masses in each spectrum. The approach is demonstrated on biological tissue datasets acquired using a DESI-MSI source interfaced with time-of-flight (TOF) or Orbitrap ion analyzers, in both positive (ES+) and negative (ES-) ion mode.

For simplicity, we will refer to *m/z* values as “masses” throughout the manuscript.

## 3 DESI-MSI datasets

### 3.1 In-house DESI-MSI datasets

DESI-MSI data of six tissue sections from mouse (mus musculus) brain and pig (sus domesticus) liver were acquired using a Waters Xevo G2-XS QToF mass spectrometer.

Three brains of a C57BL/6 mouse model were purchased from Charles River Laboratories, while pig liver samples were obtained from a local supermarket. We used the sample preparation procedure and DESI parameters reported in Tillner et al.^20^ The mouse brain samples were scanned at a rate of 75 µm/s horizontally, while the pig liver sample was scanned at a rate of 100 µm/s.

The RAW spectra from the six acquired TOF DESI-MSI were first converted into imzML using the MassLynx SDK (v4.7.0) and Python’s pyimzML package*. Then, they were filtered from the baseline noise using a modified version of the *kneedle* algorithm^21^ (Section S1 of Supplementary Information) and smoothed using a Savitzky-Golay kernel convolution. Centroided peaks were detected and used as input for the recalibration procedure (Section S2 of Supplementary Information). Throughout the manuscript, we will refer to spectrum and list of centroid peaks as synonyms.

### 3.2 Public DESI-MSI datasets

Twenty-four DESI-MSI datasets were downloaded from the public service METASPACE^†^. We considered various tissue types analyzed with either Orbitrap or TOF ion analyzers. Details about the datasets can be found in Table S1 of Supplementary Information.

The datasets consisted of centroided RAW peaks. No peak detection or denoising was applied.

## 4 Methods

### 4.1 Selection of sample pixels

The first step of the presented workflow consists of determining the pixels associated with the biological sample. Since our method uses endogenous molecules that are expected to be detected by DESI-MS in tissue, it is crucial to remove all pixels that do not correspond to tissue-related signals.

The procedure aims at discriminating the sample-related from off-sample pixels, using a supervised classifier trained on a set of user-defined labeled pixels.

We first define the set of features from the RAW peaks, applying a uniform binning with a bin size of 1 *m/z*. Subsequently, through a graphical user interface (GUI), the user manually labels a set of pixels as either ‘sample’ or ‘background’ (Figure S2 of Supplementary Information). Then, a linear support vector machines (SVM) model is fitted on this set of features and labels and used to predict all pixels’ labels, providing a binary map of the region-of-interest (ROI). The user can manually refine the segmentation through the GUI. Finally, connected regions (8-neighborhood) smaller than a given size are assigned to the background class. In all the experiments, we set the size threshold equal to 50 pixels.

The ROI mask is saved in a comma-separated values (CSV) file to be used in the workflow’s following steps.

### 4.2 General recalibration workflow

In this section, a general description of the recalibration method is given, with details reported in the following sections.

Let us consider a DESI-MSI dataset representing a collection of *N* sequentially acquired spectra (where *N = N*^(ROI)^ +*N*^(off)^) is the sum of the number of sample ROI and off-sample pixels).

The calibration procedure aims to estimate the exact (calibrated) mass from the observed mass of the detected ions.

One common strategy is based on reference points, called *lock masses*.^22^ This method assumes a statistical model *g* between the exact **µ** = (*µ*_*k*_) and observed 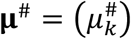 masses:

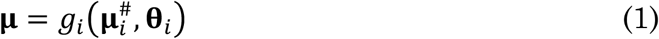

where the index *i* indicates that the observed masses **µ**^#^, the model *g* and its parameters **θ** are spectrum (or pixel)-specific.

Given a set of reference masses and their observed values, the parameters are fitted from data, and the model is used to predict the calibrated values of all observed masses.

When using endogenous signals as a reference, lock masses can be spectrum-specific^19^ (**µ** = **µ**_*i*_) or be defined globally. Our method uses the latter approach.

When using a global set of reference masses, recalibration model can only be fitted if **µ**^#^ are available in all pixels. However, due to the molecular heterogeneity of biological samples, this property is not guaranteed. To overcome this challenge, we model the observed **µ**^#^ as a smooth function of the acquisition time (or, equivalently, the pixel order), which is equivalent to assuming that the masses **µ**^#^ depend on the actual conditions of the instrument, and that these smoothly vary in a controlled environment. In practice, we model each mass in **µ**^#^ as a time series, where the pixel order is a proxy variable for the acquisition time.

Given a reference mass *M* ∈ **µ**, let **P**_*M*_ = (*p*_1_, *p*_2_, …) = (*p*_*i*_)be the subset of ROI pixels where it is detected with mass 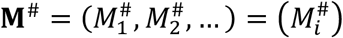, where 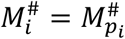. We model the time series trend using a generalized additive model (GAM) ^23^:

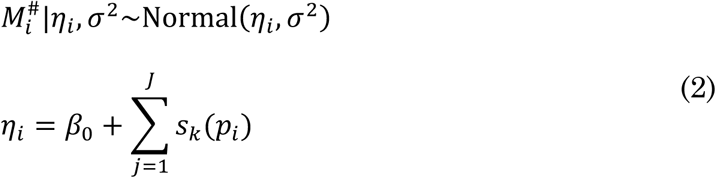

where *s*_*l*_ are penalized cubic spline functions and *J* is equal to 20.

The procedure, repeated for all *M* ∈ **µ**, separately, provides the models for all reference masses. As for any regression models, if **P**_*M*_ corresponds to ‘a large portion’ of the ROI, the fitted model can accurately predict the values of 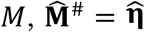, that would have been observed (up to a random error term) in all ROI pixels, subject to the instrumental condition at the acquisition time.

In the following section, we describe the procedure to detect and select the reference ions signals.

### 4.3 Detection and selection of reference ions

Since the molecular composition of the sample is largely unknown, it is practically impossible to define a general set of lock masses that works for all datasets. Before describing the procedure aimed at identifying them, we need to introduce properties that define ‘good’ endogenous reference masses.

The candidate reference mass is optimal if: a) it is detected in a ‘large’ portion of the sample, b) its peak is ‘isolated’ in the *m/z* space. Property (a) allows us to use the model described in Eq. 2, while property (b) reduces the uncertainty of the matching procedure.

As the reference ions consist of molecules that are expected to be found in tissue samples and be detected by DESI-MS, we perform a database search to determine the list of candidates. The database **Ω** includes phospholipids, fatty acids, mono/di/triglycerides, cholesterols, and ceramides from Lipidmaps^24^ and Human Metabolite DataBase^25^ (HMDB) databases. Deprotonated and chlorine adducts ([M-H]^-^, [M+Cl]^-^) are considered for negative polarity mode, while protonated, sodium and potassium adducts ([M+H]^+^, [M+Na]^+^, [M+K]^+^) are considered for positive polarity mode.

Given a database mass *M* ∈ **Ω**, a mass *m*_*i*_ among observed masses in pixel *p*_*i*_ is considered a candidate match if *m*_*i*_ ∈ [*M* − λ, *M* + λ], where λ = *W* × *M* × 10^,-^ is the mass distance from *M* corresponding to the relative error *W* in ppm units. All masses satisfying the condition are considered possible matches, meaning that more matches per pixel can be found. For each candidate match, we retrieve its peak index *π*_*i*_, observed mass 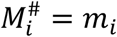, and intensity 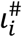.

To satisfy property (a), for all *M* ∈ **Ω**, we remove the candidate matched peaks that are detected in less than 75% of the sample ROI pixels.

Subsequently, we further filter the list of candidate reference ions using a method based on a kernel density estimator (KDE) following a similar procedure to that described in Smirnov et al.^26^ If candidate reference peaks for *M* are represented as points with coordinates 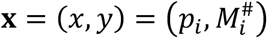, we assume that highly dense connected regions represent the shift trends of *M*.

Given a candidate reference mass *M* ∈ **µ** ⊆ **Ω** and its observed values 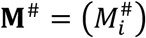, we estimate the density 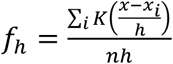 of the (*x, y*)-points using a 2D Fast Fourier

Transform (FFT) KDE ^27^, with a triangular kernel *K*, defined as

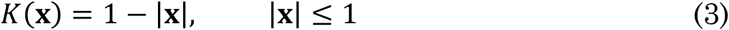

The kernel is fitted on the coordinates of the points scaled to [0,1] interval.

The triangular kernel is chosen because of its computational efficiency. The kernel bandwidth is set to *h* = 2.576 × *σ* × *N*^−1/5^, where *σ* represents the standard deviation of the whole set of points and *N* represents the number of points **x**.^28^ The kernel is fitted on a regular grid of size *G* × *G*, with *G* = 1024.

Given the estimated 2D density 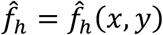, we identify the curve passing through its local maxima as follows. For each *x*′ ∈ {1, …, *G*}, we determine the local density maximum 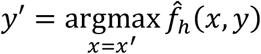. If 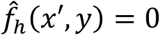 for all *y*, no maximum is considered for that value of *x*′. Subsequently, a cubic spline *S*(*x*) is fitted on the vector of local maxima 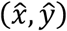 after transforming them back to the original (pixel, mass)-space. Using the predicted spline values *S*(*p*_*i*_) for all *p*_*i*_, we calculate the absolute residuals *r*_*i*_ and the dispersion *d*_*i*_:

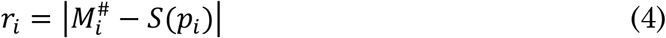

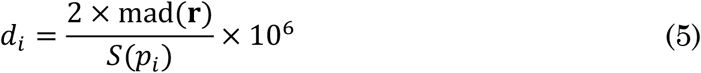

Points with *r*_*i*_ ≥ 2 × mad(**r**), **r** = (*r*_*i*_), where “*mad*” is the median absolute deviation multiplied by the inverse of cumulative Normal distribution 1/Φ^−1^(3/4) ≈ 1.4826, are considered outliers and removed from the list of matched peaks (Figure 1A). Finally, the candidate reference *M* is kept if: the number of distinct inlier pixels is greater or equal than 75% of the ROI size and 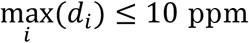. In this way, we select isolated signals (property (b)).

**Figure 1.**
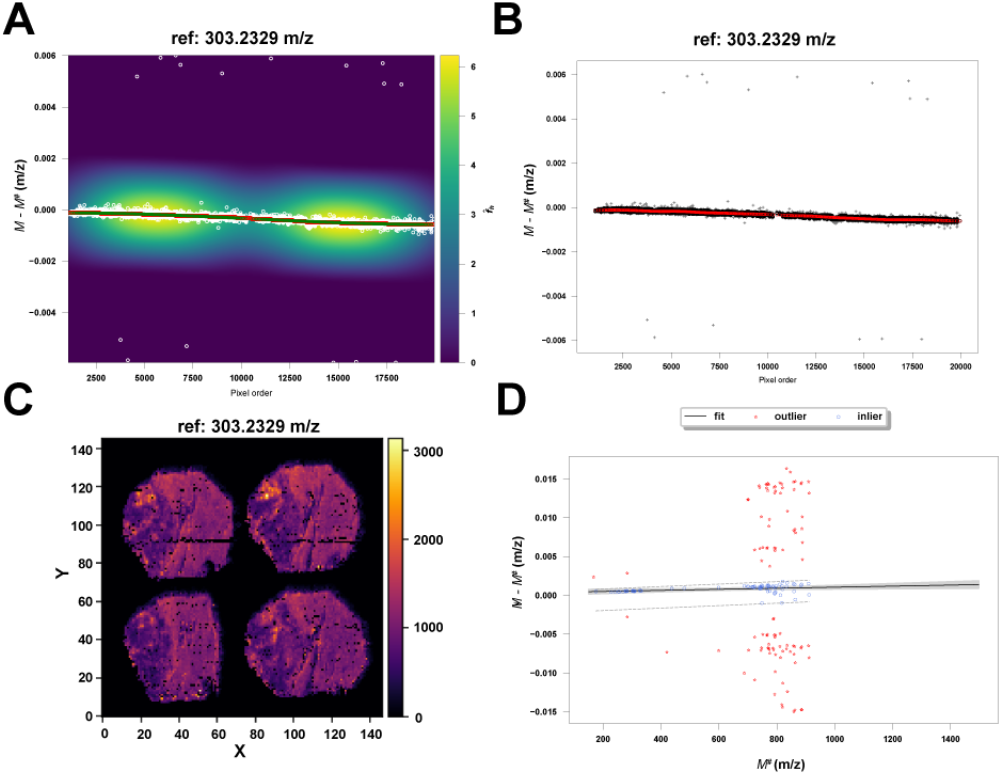
Example of the recalibration process for the Orbitrap dataset liver ES-. (A) FFT KDE is fitted from the matched masses for a candidate reference (303.2329 m/z). The white points represent the difference in m/z between the observed and the theoretical values of the reference. The curve passing through the density maxima (points in red) is used to fit a regression spline (blue points). (B) Points with large residuals from the spline are labeled as outliers (in grey) and excluded from the matched set. The remaining inliers are then used to fit a GAM (red line). (C) The intensity corresponding to the inlier points can be plotted to reveal their spatial distribution. This allows validating the consistency of the selected matches visually. (D) Finally, a regression model is fitted in each pixel using the only reference masses with similar errors in ppm (blue circles) within the residual intervals defined by the filter (grey dashed lines) (Eq. 6-7). The degree of the polynomial corresponds to the smallest BIC value. The predicted values (black line, the grey bands represent 95% confidence intervals) are used to correct all observed masses in the pixel.

The procedure is repeated for all candidate reference masses.

The final set of reference masses is denoted as **µ**^∗^. The time series trends (Eq. 2) for each *M*^∗^ ∈ **µ**^∗^ are fitted using the corresponding inlier points, and the models are used to predict the observed values 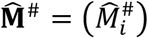 in all ROI pixels (Figure 1B). The intensities 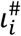 of the matched peaks can be plotted to reveal their spatial distribution (Figure 1C). Visual inspection provides additional validation of the consistency of the matched peaks. When multiple matches are available per pixel, the intensities of the peaks with the smallest *r*_*i*_ are plotted.

### 4.4 Pixel-wise mass recalibration

Although the selected reference ions have passed the filtering procedure, mismatched peaks may still be present if: a) same peaks fall within two search windows so that they are assigned to two reference ions, b) shifted peaks fall within the search window by chance (especially if the mass shift is large).

To reduce the chance of using mismatched reference ions masses, we apply a pixel-specific mass filter, based on the assumption that most of the true matches share a similar relative mass error. The details of the filtering procedure are reported in Section S4 of Supplementary Information.

Once the set of reference masses 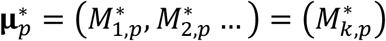 is determined for pixel *p* together with the values predicted by GAMs, we fit a model *g* (Eq. 1) specific for each type of ion analyzer (Suppl. Information S3). For Orbitrap analyzers, we fit a linear regression model ^29^

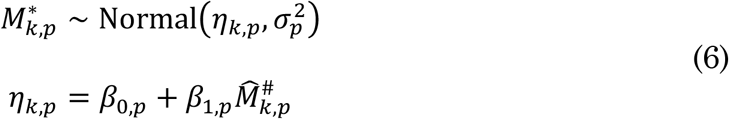

Instead, for the TOF analyzer, we use a polynomial regression model ^30,31^

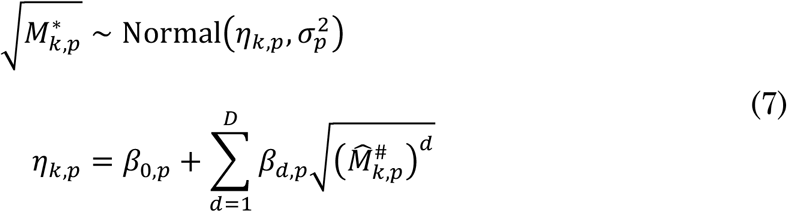

where the optimal polynomial degree *D, D* ≤ 5, corresponds to the smallest Bayesian Information Criterion (BIC).^32^

The fitted model is then used to predict the calibrated mass for all the detected ions in the specific pixel (Figure 1D).

## 5 Results and discussion

As a first experiment, we tested the validity of the pixel-wise recalibration model and the reference mass filtering described in Section S4 of Supplementary Information. We simulated a series of pixel-wise reference masses and their observed values following the method described in Section S5 of Supplementary Information (Figure S1 of Supplementary Information). In all simulations, we observed a significant reduction of the median relative error, in both Orbitrap andTOF models. No significant difference in terms of performances was observed between the two models. Also, the simulations confirmed a reduction of the heteroskedasticity of the selected masses (Figure S5 of Supplementary Information).

We then tested the DESI-MSI datasets after removing the pixels outside of the ROI as described in Section 4.1.

All recalibration parameters were kept fixed within the Orbitrap or TOF datasets, as described in the Methods section. For Orbitrap datasets, we used a search window *W* = 20 ppm, while for TOF datasets we used *W* = 100 ppm.

FFT KDEs for matched points were fitted using the “KDEpy” package for Python‡ (v. 1.1.0). The cubic splines are fitted using the function *UnivariateSpline* available in “SciPy” package for Python (v 1.7.1).^33^ The smoothing parameter *s* of the function was determined by 5-fold cross-validation, among 20 values from 0.0001 to 0.1 evenly spaced in the log_10_ scale, corresponding to the smallest mean squared error.

The KDE-based model of the reference mass shift showed its robustness in the case of close peaks, correctly discarding wrongly matched peaks (Figure 2).

**Figure 2.**
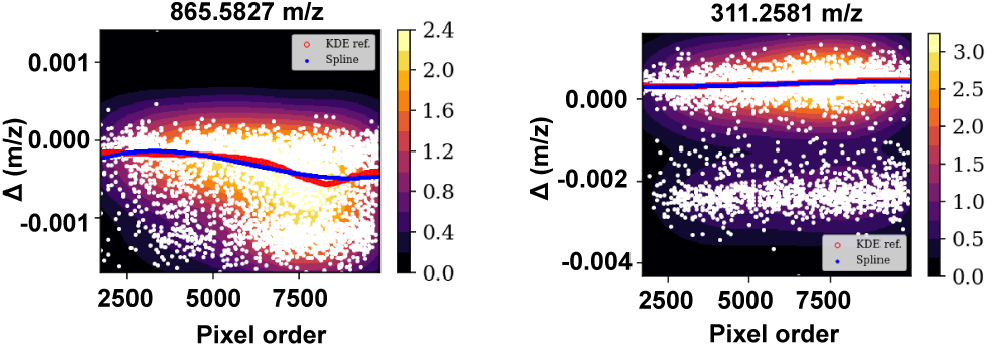
Left: Two peaks are too close to be resolved by the KDE. This reference is correctly discarded because of the large dispersion around the regression spline. Right: in this case, the KDE can resolve the two close peaks. Here, the method correctly fits the spline on the points belonging to one of the peaks. After filtering the outliers, the dispersion is below the threshold.

The GAMs were fitted using the pyGAM package (v. 0.8.0) for Python.^34^ The spline penalization parameter was chosen among eleven values varying between 0.001 and 1000 evenly spaced in the log_10_ scale, corresponding to the smallest generalized cross-validation value (GCV).^35^

In all tested DESI-MSI datasets, except for Orbitrap pancreas ES-, the selected reference masses for the pixel-wise recalibration well covered their acquisition *m/z* range (Figure S3-S4 of Supplementary Information). In general, the reference masses were more concentrated in the *m/z* ranges below 400 and above 600, corresponding to small molecules and phospholipids, respectively.

In five TOF datasets, the optimal pixel-wise recalibration models were polynomial with a degree greater or equal to two. In particular, the distribution of the regression models coefficients revealed that the TOF mass shifts were greater than Orbitrap (Orbitrap error within 5 ppm, while TOF error up to 65 ppm) (Figure 3). This observation may indicate a higher tendency of one class of analyzers to be subject to the changes of the environmental conditions, although these may depend on the unknown actual instrumental conditions for each dataset.

**Figure 3.**
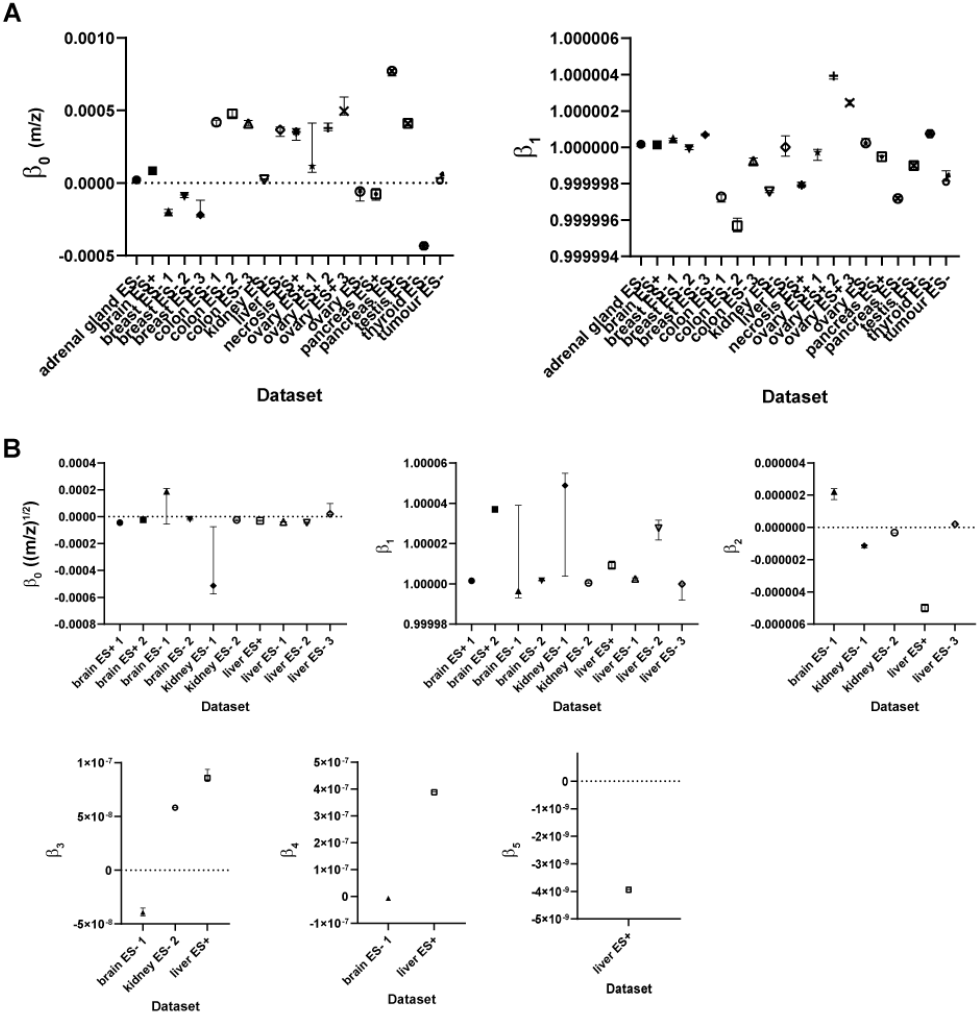
Distribution of the pixel-wise recalibration regression model coefficients. Each dataset is represented by the median and 0.95 quantile interval of the model coefficient values. Intercept and slope of Orbitrap linear models are shown in (A). In five TOF datasets (B), the models were polynomial with a degree greater or equal to two. In all cases, the linear coefficient β_1_ was significantly larger than the intercept β_0_, with β_2_ M ≫ β_0_.

To evaluate the recalibration efficacy, we looked at two main effects: 1) the number of putative molecular annotations, 2) mass error of test ions.

The number of putative annotations was generated using the METASPACE website. METASPACE assigns molecule identities to MSI datasets based on a metabolite-signal match score (MSM) calculated from spectral and spatial measures and an FDR-estimation using a decoy strategy. Each assignment is characterized by an FDR equal to 5%, 10%, 20%, and 50%. The annotation is performed by database search. Because of the biological nature of the analyzed samples, we selected the following four databases among those available: a) “ChEBI 2018-01”, b) “CoreMetabolome v3”, c) “HMDB v4”, d) “LipidMaps 2017-12-12”.

All METASPACE default parameters were used. Specifically, the interval for the image generation was set to 3 ppm, and the decoy set size was set to 20. We only considered the number of unique annotations corresponding to the stringent criterion of FDR = 5%. In addition, we considered distinct annotations corresponding to different molecular formulas plus their adduct among the four used databases. This choice was justified by the fact that the actual molecules may be present in only specific databases. The annotations were performed using the ROI filtered raw and recalibrated peaks without applying any peak binning or spatial filter.

We checked whether the largest number of unique annotations was found in the original or recalibrated dataset. Also, we calculated the relative difference between the number of annotations in the two datasets:

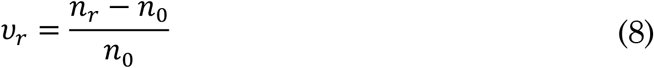

where *n*_0_ is the number of annotations in the original dataset, and *n*_*r*_ in the recalibrated dataset.

In twenty-two cases (≈ 73%), the recalibrated dataset received a larger number of assignments compared to the raw dataset (Figure 4).

**Figure 4.**
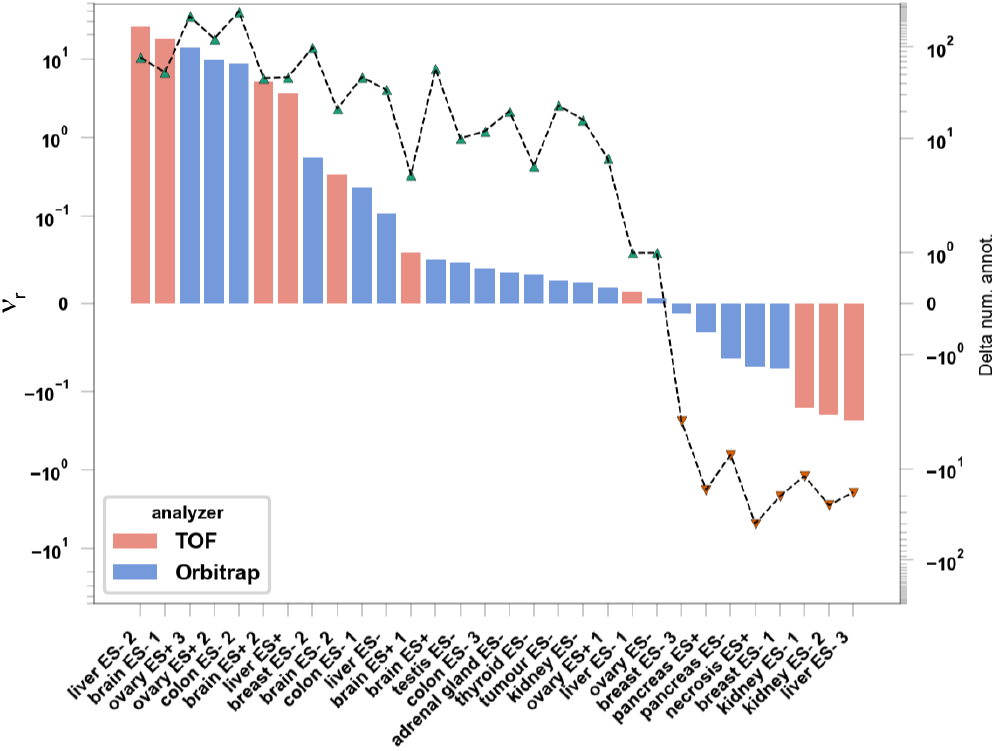
Results of the METASPACE annotations with an FDR=0.05. The bars (left axis) represent the relative difference of the number of annotated molecules between the original and recalibrated datasets. The bars corresponding to TOF datasets are colored red, while those corresponding to Orbitrap datasets are colored in blue. The dashed line and triangular markers (right axis) represent the actual difference between the number of annotated molecules in the original and recalibrated datasets (Eq. 8). The triangles are colored in green if the recalibrated data generated more annotations than the original data, in red otherwise. In eight cases (five Orbitrap and three TOF), the recalibrated data generated fewer annotations.

The recalibration increased the number of annotations by up to 246. In contrast, in TOF liver ES-3 we observed the largest relative decrease equal to 23% of the original annotations, corresponding to 18 fewer annotations, while the largest absolute decrease was observed in Orbitrap necrosis ES+ with 40 fewer annotations, equivalent to about 7% of the original annotations (Figure 4).

We tested the mass accuracy of a set of test masses to evaluate the effect of the recalibration. We followed the same idea described in Boskamp et al.^18^ Since the actual molecular content of a sample is unknown, we compared the mass accuracy of a list of ions expected to be detected by DESI-MSI in biological tissue samples.

The test ions list was generated from the available annotated molecules in the public datasets of METASPACE. The list consisted of the monoisotopic form plus the first three identified isotopes detected in more than 10% of the datasets from the same tissue type and ion mode of the DESI-MSI dataset. We used a Python script from LaRocca et al.§ (version available in June 2021) to generate the list of candidate test masses. The mass values common to the set used for fitting the pixel-wise recalibration models were excluded. In all datasets, the test masses covered the acquisition *m/z* range, confirming that they represented a good set of probes for evaluating the recalibration effects (Figure S3-S4 of Supplementary Information).

Using the procedure described in Section S6 of Supplementary Information, we selected the test masses within 1 ppm from the most abundant relative errors and calculated the median of the pixel-wise median difference between the absolute mass errors 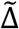. We tested if we could reject the null hypothesis 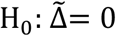, using bootstrapping (number of repetitions equal to 10^4^). We observed in nineteen datasets a significant (Benjamini-Hochberg corrected p-value < 0.05) decrease of the median relative error after the recalibration, with values varying between 0.02 and 63.6 ppm. In all these datasets, the final median relative error was below 2.7 ppm for Orbitrap, and below 4.9 ppm for TOF. In one case (TOF liver ES-3), the median relative error increased by about 1.3 ppm, with a final median relative error equal to 4 ppm. In ten datasets, 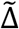 was not significantly different from zero (Figure 5, Table S2-S3 of Supplementary Information).

**Figure 5.**
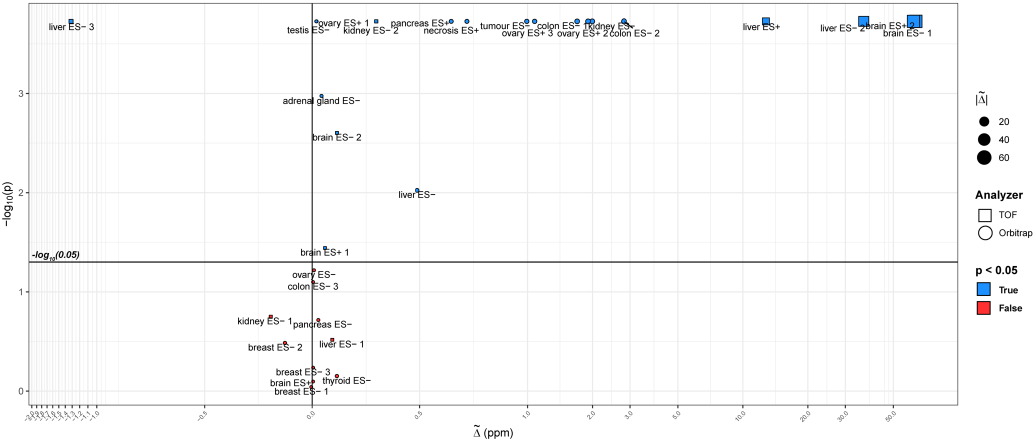
Scatter plot representing the bootstrapping results for the median difference between the mass errors in the original and recalibrated datasets. Each dataset is characterized by its 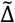 value (calculated following the method described in Section S6 of Supplementary Information) and the significance value of the bootstrapping test. Most datasets show a significant reduction of the test masses absolute error, with values up to about 60 ppm.

Consistently, the datasets with the largest increase in the number of annotations corresponded to those with the largest decrease in the mass error.

Furthermore, the recalibration improved the ion image quality. After removing the mass shifts, all pixels correctly displayed the expected molecular spatial distributions (Figure 6).

**Figure 6.**
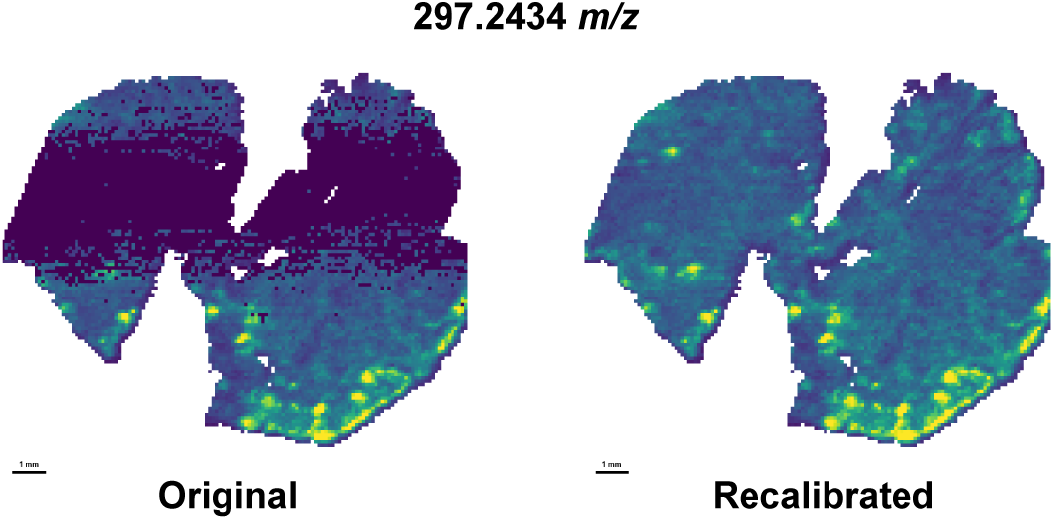
Example of the effect of recalibration on the quality of an assigned metabolites’ spatial distribution by METASPACE for the ORBITRAP colon ES-2. The original dataset shows a band of missing intensities caused by the masses’ nonlinear distribution across the pixels, which is corrected in the recalibrated data.

Finally, we tested the effect of modeling the reference mass shifts as time series. We repeated the pixel-wise recalibration, considering the pixels’ spectra independent. We searched the candidate reference using the same method described in Section 4.3, with a tolerance *W* = 20 ppm for Orbitrap, and *W* = 100 ppm for TOF datasets. The masses that were matched in less than 10 pixels were removed. Thus, in each pixel we applied the pixel-wise recalibration procedure described in Section 4.4. Subsequently, we tested the mass accuracies for the METASPACE masses, using the procedure described in Section S6 of Supplementary Information. Finally, we tested the null hypothesis 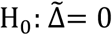, using bootstrapping (number of repetitions equal to 10^4^). The results showed that in fifteen datasets there was a significantly (Benjamini-Hochberg corrected p-value < 0.05) smaller relative error when the references were modeled as a time series. In eleven cases, the relative error was smaller in the model with independent pixel-wise regressions. In nine datasets, we did not observe a significant difference (Figure S6 of Supplementary Information). However, the values of 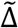 varied between 0.004 and 0.2 ppm in the cases favorable to the independent models, while they varied between 0.03 and 35.2 ppm for the results favorable to the time series models. These results suggested that modeling the reference masses as a time series was more beneficial than considering independent models for each pixel spectrum.

## 6 Conclusion

Despite its enormous potential to capture the spatial characteristics of the metabolic and proteomic mechanisms in a wide range of biological samples, MSI remains a relatively young technology. Advancements in fundamental data quality control are necessary to transform it into a more reliable approach.

Accuracy of measured masses is particularly challenging in imaging approaches since usually a run consists of tens of thousands of pixels and may require several hours to complete. Thus, the changes of the instrumental condition during this time may invalidate its initial configuration, resulting in mass drifts that correlate with time or pixel order.

Here, we have presented a computational workflow that exploits the presence of typical biological molecules in most of the sample spectra and uses them as a set of reference ions for applying a spectra-wise lock mass correction.

Using a global set of reference ions for the entire MSI dataset allowed us to test the assignment’s validity by visualizing the corresponding spatial distributions. Furthermore, we modeled the mass drifts as a smooth function of the acquisition time, as expected in acquisitions occurring in a controlled environment. Thus, these models represent an additional level of evidence about the assignments’ correctness.

FFT KDE-based reference match filtering, together with GAMs, proved robust against outliers and false-positive reference ions, for instance, in the presence of close peaks with a complementary spatial distribution to that of the actual reference ions peaks.

We showed that the presented approach could improve the mass accuracy, removing the nonlinear fluctuations of the measured masses. Additionally, our approach improved the molecular assignment to the detected peaks, using state-of-the-art molecular annotation methods, such as METASPACE, and their quality in terms of assigned spatial distributions.

Modeling the drifts of the reference masses as a time series resulted in better performances compared to considering the pixels’ spectra independent.

In the dataset with already accurate masses, the recalibration can introduce errors due to its statistical nature. For this reason, it is crucial to evaluate the test masses accuracies before and after performing the recalibration, using the methods described here.

We employed pixel-wise recalibration models specific to the physics of the MS analyzer. This is an essential aspect of the procedure since these characteristics influence the statistical properties of the signal generated from the detected ions. It is also important to underline that we designed the mass shift models to be simple, considering only the time-dependency of the acquisition. Unfortunately, this means that several complex aspects of the phenomena driving the mass drifts are not captured. This is a necessary trade-off between generality and reduction of the risk of overfitting. More realistic models may be designed if additional variables are tracked, such as temperature, humidity, just to name a few.

We showed that computational models could effectively reduce the mass drifts that affect MSI datasets. However, it is crucial to stress that they can represent a temporary solution to this problem until technological solutions capable of performing an accurate online calibration will not be available.

The presented work has limitations. Although we used an extensive list of publicly available molecular masses as references, some peculiar molecules for unknown samples may be missing. For this reason, building a database of accurately identified molecules in previous studies is crucial.

Another limitation of the approach is that it can only be applied to the sample-related ROI pixels. Thus, when selecting these pixels, the spatial information necessary to identify possible non-sample-related signals is lost. However, this difficulty can be easily overcome by applying spatial-aware filters, such as SPUTNIK ^36^, before the recalibration. In this way, most of the signal correlated with regions outside of the ROI is removed before performing the pixels selection. This aspect is crucial for the correct biological interpretation of the observed molecular spatial patterns.

In the future, we will work to extend the method to other types of ion analyzers and will study the possible integration with the usage of external lock mass reference ions.

## 7 Author’ Contributions

P.I. designed the model and the computational framework and analyzed the data. H.X.H. and V.W. performed the DESI-MSI experiments. M.R.L. and Z.T. conceived the study and oversaw overall direction and planning. All authors discussed the results and contributed to the final manuscript. All authors reviewed the manuscript.

## 8 Code availability

The Python code is freely distributed under the MIT license and can be downloaded from https://github.com/paoloinglese/DESI_MSI_recal, together with the reference masses databases.

## 11 Ethics approval and consent to participate

Three snap-frozen brains of a C57BL/6 mouse model were purchased from Charles River Laboratories. No experiment was carried out on live animals.

## 12 Consent to publish

Not applicable.

## 13 Acknowledgments

We thank Clara L Feider, Meghan P. Friis, Kyana Garza, Emrys Jones, James McKenzie, Nicole Strittmatter, Jialing Zhang, for sharing their datasets on METASPACE.

## Footnotes

* https://github.com/alexandrovteam/pyimzML

† https://metaspace2020.eu/

‡ https://github.com/tommyod/KDEpy

§ https://github.com/LaRoccaRaphael/MSI_recalibration/blob/master/recalibration/internal_calibrants_generation.ipynb

