## Supplementary Information for "Mass recalibration for desorption electrospray ionization mass spectrometry imaging using endogenous reference ions"

**[1]National Phenome Centre, Imperial College London, Hammersmith Campus, IRDB Building, London W12 0NN, United Kingdom**

**[2]Section of Bioanalytical Chemistry, Division of Systems Medicine, Department of Metabolism, Digestion and Reproduction, Imperial College London, South Kensington Campus, London SW7 2AZ, United Kingdom**

### Supplementary Information

| ORBITRAP | | | | | | |
|---|---|---|---|---|---|---|
| **Uploader** | **Organism** | **Tissue** | **Analyzer** | **Ion mode** | **MS Resolution** | **METASPACE** |
| Nicole Strittmatter | Mus musculus | Adrenal gland | Orbitrap | ES- | 140k @200 m/z | YES |
| Nicole Strittmatter | Rat norvegicus | Brain | Orbitrap | ES+ | 140k @200 m/z | YES |
| Kyana Garza | Homo sapiens | Breast | Orbitrap | ES- | 60k @200 m/z | YES |
| Kyana Garza | Homo sapiens | Breast | Orbitrap | ES- | 60k @200 m/z | YES |
| Kyana Garza | Homo sapiens | Breast | Orbitrap | ES- | 60k @200 m/z | YES |
| James McKenzie | Homo sapiens | Colon | Orbitrap | ES- | 70k @200 m/z | YES |
| James McKenzie | Homo sapiens | Colon | Orbitrap | ES- | 70k @200 m/z | YES |
| James McKenzie | Homo sapiens | Colon | Orbitrap | ES- | 70k @200 m/z | YES |
| Nicole Strittmatter | Rat norvegicus | Kidney | Orbitrap | ES- | 140k @200 m/z | YES |
| James McKenzie | Homo sapiens | Liver | Orbitrap | ES- | 100k @200 m/z | YES |
| Meghan P. Friis | Homo sapiens | Necrosis | Orbitrap | ES+ | 140k @200 m/z | YES |
| Clara L Feider | Homo sapiens | Ovary | Orbitrap | ES- | 60k @400 m/z | YES |
| James McKenzie | Homo sapiens | Ovary | Orbitrap | ES+ | 100k @200 m/z | YES |
| James McKenzie | Homo sapiens | Ovary | Orbitrap | ES+ | 100k @200 m/z | YES |
| James McKenzie | Homo sapiens | Ovary | Orbitrap | ES+ | 100k @200 m/z | YES |
| Meghan P. Friis | Homo sapiens | Pancreas | Orbitrap | ES- | 140k @200 m/z | YES |
| Meghan P. Friis | Homo sapiens | Pancreas | Orbitrap | ES+ | 140k @200 m/z | YES |
| Nicole Strittmatter | Rat norvegicus | Testis | Orbitrap | ES- | 140k @200 m/z | YES |
| Jialing Zhang | Homo sapiens | Thyroid | Orbitrap | ES- | 60k @400 m/z | YES |
| Nicole Strittmatter | Mus musculus | Tumor | Orbitrap | ES- | 140k @200 m/z | YES |
| **TOF** | | | | | | |
| **Uploader** | **Organism** | **Tissue** | **Analyzer** | **Ion mode** | **MS Resolution** | **METASPACE** |
| - | Mus musculus | Brain | Xevo G2-XS qTOF | ES- | 20k @200 m/z | NO |
| Emrys Jones | Mus musculus | Brain | TOF reflector | ES- | 30k @585 m/z | YES |
| - | Mus musculus | Brain | Xevo G2-XS qTOF | ES+ | 20k @200 m/z | NO |
| - | Mus musculus | Brain | Xevo G2-XS qTOF | ES+ | 20k @200 m/z | NO |
| Emrys Jones | Mus musculus | Kidney | qTOF | ES- | 30k @800 m/z | YES |
| Emrys Jones | Sus domesticus | Kidney | TOF reflector | ES- | 30k @585 m/z | YES |
| - | Sus domesticus | Liver | Xevo G2-XS qTOF | ES- | 20k @200 m/z | NO |
| - | Sus domesticus | Liver | Xevo G2-XS qTOF | ES- | 20k @200 m/z | NO |
| Emrys Jones | Sus domesticus | Liver | TOF reflector | ES- | 30k @500 m/z | YES |
| - | Sus domesticus | Liver | Xevo G2-XS qTOF | ES+ | 20k @200 m/z | NO |

*Table S 1 - Details of thirty tested DESI-MSI datasets. The "MS Resolution" column represents the approximate mass resolution at a given m/z value. The "METASPACE" column reports the datasets that were downloaded from the METASPACE website (YES) and those that were acquired in our experiments (NO).*

#### S.1 TOF-MS RAW denoising

A two-step procedure was applied to remove baseline (BN) and signal noise (SN) independently. First, the BN level is identified using a modified version of Zhurov et al.[1] As in the original method, we assume that, in a mass spectrum, the signal intensities are marginally distributed as a mixture of two components: a BN component and a sample signal component (with its noise, SN). The BN filtering's main scope identifies the optimal point of separation between two distributions (the "cut level") such that only the sample signal is preserved. To avoid the ambiguity introduced by selecting the number of histogram bins used in Zhurov et al., we use a different approach to identify the cut level. All signal intensities are sorted in decreasing order, and the cut point is chosen at the *elbow point* of the curve. This choice assumes that the noise level represents the intensity at which the most noticeable intensity distribution change is observed (maximum curvature point). Since noise can affect the estimated value of the second derivative, we identify the elbow point using a simplified version of the *Kneedle* algorithm[2] that only looks at the farthest point from the diagonal line connecting the two extremal points of the curve.

All zeros are removed from the signal, and, as described in Zhurov et al., signal intensities are log-transformed before the BN level estimation. The elbow point is used as a cut-off, and all signal intensities below that value are set to zero.

The second component of the noise, SN, is assumed to be signal-dependent (multiplicative noise). We use a smoothing procedure based on a local transformation of the signal through Savitzky-Golay convolution to remove SN. All negative intensities generated by the application of the Savitzy-Golay filter are set to zero.

The Savitzky-Golay filters were estimated with a degree equal to three and a half-window equal to four points in the presented experiments.

Smoothing followed by peak detection was preferred to one-step approaches, such as wavelet peak-picking[3], for computational efficiency reasons.

#### S.2 Peak detection

A denoised signal (as described in the previous section) was used to determine peak mass-to-charge values and intensities.

Peaks and base points (half peak intensity) were detected using the function *find_peaks* available in Python's Scipy library.[4] Minimum width of 3 points and a minimum prominence equal to half of the BN value were used to retain the signal peaks. The centroid *m/z* value calculated within the base points interval was assigned to the peaks.

Since the smoothing modifies the original intensities, we assigned each peak the maximum RAW intensity in the interval between the peak base points. This ensures that the peak intensity is consistent with the original RAW data.

#### S.3 Recalibration models for TOF and Orbitrap ion analyzers

Recalibration regression models for TOF and Orbitrap MS can be derived from the equations relating the measured quantities and the *m/z* values.

In the case of the TOF analyzer, the relationship between the observed time-of-flight $t$ and the mass $m$ and charge $q$ of an ion is given by the formula:

$$\sqrt{\frac{m}{q}} = \beta_1 t + \beta_0 \qquad \text{(S1)}$$

where the coefficients $\beta$ represent the instrumental condition and geometry.

Lock mass correction is performed by applying Eq. S1 to a set of one or more reference ions with known mass $m$ and estimate the coefficients $\beta$:

$$\sqrt{\left(\frac{m}{q}\right)_{i,t}} = \hat{\beta}_1 t_{i,o} + \hat{\beta}_0 = \hat{\beta}_1{}' \sqrt{\left(\frac{m}{q}\right)_{i,o}} + \hat{\beta}_0{}', \qquad i = 1, \dots, n \qquad \text{(S2)}$$

where $n$ is the number of lock masses, and the indices $t$ and $o$ represent the nominal and observed values, respectively.

Eq. S2 can be extended to a polynomial of degree $d$ in the square-root of $m/q$:[5,6]

$$\sqrt{\left(\frac{m}{q}\right)_{i,t}} = \beta_0 + \sum_{p=1}^{d} \beta_p \sqrt{\left(\frac{m}{q}\right)_{i,o}^{p}} \tag{S3}$$

The calibration coefficients can be estimated by fitting a polynomial regression model on the lock masses. Thus, the coefficients $\boldsymbol{\beta}$ can be used to estimate the theoretical (calibrated) mass for all observed masses using Eq. S3.

An analogous procedure can be used to determine the calibration equation for Orbitrap.

In the case of the Orbitrap analyzer, the measured frequency $f$ is related to the mass $m$ and charge $q$ of an ion by the equation:

$$\frac{m}{q} = \frac{\beta_1}{f^2} + \beta_0 \tag{S4}$$

where $\beta$ are the calibration coefficients.[7]

By substituting the observed frequency with the corresponding observed $m/q$, the lock mass equation becomes linear in $m/q$.[8] Given $n$ reference ions:

$$\left(\frac{m}{q}\right)_{i,t} = \hat{\beta}_1 \left(\frac{m}{q}\right)_{i,o} + \hat{\beta}_0, \qquad i = 1, \dots, n \tag{S5}$$

The calibration coefficients $\hat{\beta}$ can be estimated by fitting a linear regression model on the lock masses.

Afterward, once the coefficients $\hat{\beta}$ are estimated, it is possible to calibrate all observed masses using Eq. S5.

#### S.4 Mass filter for pixel-wise regression

Given a pixel $p$, and the candidate matched masses $\widehat{\mathbf{M}}_p^{\#}$, we assume that the relative mass errors of the true matches are distributed in a small interval.

Thus, we select the masses corresponding to an interval of mass errors around its highest density peak. First, in each ROI pixel $p$, for each reference mass $M_k$ and its predicted observed value, $\widehat{M}^{\#}_{k,p}$, we calculate the relative errors in ppm units:

$$\delta_{k,p} = \delta_p(M_k) = \frac{M_{k,p} - M_k}{M_k} \times 10^6 \tag{S6}$$

Given the relative errors of all reference masses in $p$, $\boldsymbol{\delta}_p = (\delta_{k,p})$, we estimate their density in the set of points $\mathbf{E} = \left(e_0 = \min(\boldsymbol{\delta}_p), \dots, e_j = \min(\boldsymbol{\delta}_p) + j \times 0.001, \dots, e_J = \max(\boldsymbol{\delta}_p)\right)$ using a Gaussian FFT KDE with Silverman bandwidth[9]. Therefore, we select the masses that lie in the interval $\mathbf{I} = \left[e_{(-0.75)}, e_{(+0.75)}\right] \subseteq \mathbf{E}$ around the highest peak point $e_{(\max)}$, where $e_{(-0.75)}$ and $e_{(+0.75)}$ are the left and right shoulder points corresponding to 0.75 of the $e_{(\max)}$ height, respectively.

Under the approximation of a linear relationship between the observed nominal values, given a set of observed masses $\widehat{\mathbf{M}}^{\#}_p$ with a constant relative error $\delta'$, we can write

$$\begin{aligned} \mathbf{M} &= \beta_0 + \beta_1 \widehat{\mathbf{M}}^{\#}_p \\ &\approx \beta_1 \widehat{\mathbf{M}}^{\#}_p = \frac{1}{1 - \delta' \times 10^{-6}} \times \widehat{\mathbf{M}}^{\#}_p \end{aligned} \tag{S7}$$

where we have used the fact that $\beta_0 \ll \beta_1 M^{\#}_{k,p}$, for all $M^{\#}_{k,p} \in \widehat{\mathbf{M}}^{\#}_p$.

This is equivalent to saying that the selected masses are approximately bounded by the lines with slopes $1/(1 - \min(\boldsymbol{\delta}_p) \times 10^{-6})$ and $1/(1 - \max(\boldsymbol{\delta}_p) \times 10^{-6})$ or, that the points are heteroskedastic to the line with slope equal to $1/(1 - e_{(\max)} \times 10^{-6})$ making them problematic for a regression model (Figure S1).

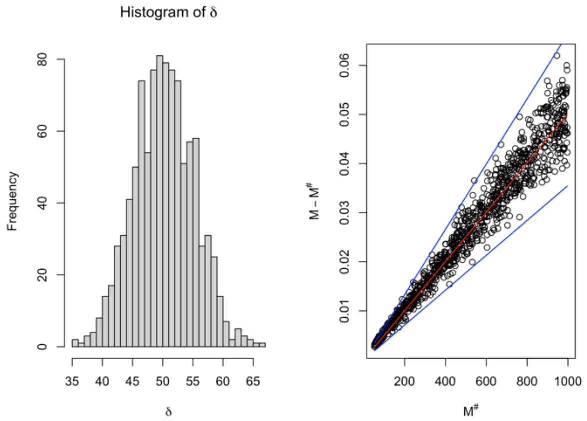


*Figure S 1 – Simulated masses following the formula* $M = 0.0005 + \beta_1 M^{\#}$*, with* $M \sim \mathrm{Unif}(50{,}1000)$*,* $\beta_1 = \delta_p M \times 10^{-6}$*, and* $\delta_p \sim \mathrm{Normal}(50, 10)$ *(left). Right: the mass errors* $M - M^{\#}$ *are bounded by the lines* $M^{\#} \times \left(\frac{1}{1-min(\delta_p)\times 10^{-6}} - 1\right)$ *and* $M^{\#} \times \left(\frac{1}{1-max(\delta_p)\times 10^{-6}} - 1\right)$ *(blue lines) around the line* $M^{\#} \times \left(\frac{1}{1-e_{(\max)}\times 10^{-6}} - 1\right)$ *(red line).*

To reduce the points' heteroskedasticity, we further filter the masses that have a similar residual to the line with slope $1/(1 - e_{(\max)} \times 10^6)$.

Given the residuals $\boldsymbol{\zeta}_p = \mathbf{M} - \widehat{\mathbf{M}}_p^{\#}/(1 - e_{(\max)} \times 10^6)$, we fit and estimate their density using a Gaussian FFT KDE with Silverman bandwidth in the points $\mathbf{E}' = \left(e'_0 = \min(\boldsymbol{\zeta}_p), \ldots, e'_l = \min(\boldsymbol{\zeta}_p) + j \times 0.00001, \ldots, e'_L = \max(\boldsymbol{\zeta}_p)\right)$. Subsequently, we select the masses that lie in the interval $\mathbf{I}' = \left[e'_{(-0.01)}, e'_{(+0.01)}\right] \subseteq \mathbf{E}'$ around the highest peak point $e'_{(\max)}$, where $e'_{(-0.01)}$ and $e'_{(+0.01)}$ are the left and right shoulder points corresponding to 0.01 of the $e'_{(\max)}$ height, respectively (Figure S 5-bottom). In this case, we retain most of the masses belonging to the highest peak to reduce the number of filtered masses. If one of the shoulder points is placed beyond another density peak, we set the interval boundary as the midpoint between the highest peak and the closest peak points. Optionally, we select a closer density curve point, such that the distance between the selected point and peak maximum point is never greater than 0.005 *m/z* (equivalent to 5 ppm for a mass-to-charge equal to 1000 *m/z*).

#### S.5 Simulation of pixel-wise recalibration

We evaluate the effect of the pixel-wise recalibration model on simulated observed and nominal masses. Given $N^{(\mathrm{true})}$ nominal masses, $\boldsymbol{m} = (m_k)$, $m_k \sim \mathrm{Unif}(50, 1000)$, we generate the observed masses $\boldsymbol{m}^{\#} = m_k^{\#}$ following a linear model in the masses (squared-root of the masses for TOF), as follows.

First, the intercept $\beta_0$ and the relative errors $\boldsymbol{\delta} = (\delta_k)$ are set:

$$\begin{aligned} \beta_0 &\sim \mathrm{Unif}(\alpha_0, \alpha_1) \\ \delta_k &\sim \mathrm{Normal}(\eta, \sigma) \end{aligned} \qquad \text{(S8)}$$

Given the values of $m_k$, $\beta_0$, and $\delta_k$, we set $\beta_1$:

$$\begin{aligned} \beta_1 &= \frac{m_k - \beta_0}{m_k(1 - \delta_k \times 10^{-6})} \qquad (\mathrm{Orbitrap}) \\ \beta_1 &= \frac{\sqrt{m_k} - \beta_0}{\sqrt{m_k(1 - \delta_k \times 10^{-6})}} \qquad (\mathrm{TOF}) \end{aligned} \qquad \text{(S9)}$$

where we have simply inverted the linear equations and used the definition of $\delta_k$ in ppm units (Eq. S6).

The observed masses $\boldsymbol{m}^{\#} = m_k^{\#}$ are then calculated by inverting the linear equation expressing the dependency of $\boldsymbol{m} = m_k$ from $m_k^{\#}$ (or the square-roots in the case of TOF) (Eq. S3-S5).

A set of $N^{(\mathrm{noise})}$ noise pairs $(m_k', m_k'^{\#})$ are added, where $m_k' \sim \mathrm{Unif}(\min(\boldsymbol{m}), \max(\boldsymbol{m}))$ and $m_k'^{\#} \sim \mathrm{Unif}(\min(\boldsymbol{m}^{\#}), \max(\boldsymbol{m}^{\#}))$.

We then apply the pixel-wise recalibration model with its filtering approach, and estimate the relative mass error of the true pairs from the predicted calibrated masses.

The simulations are run with these parameters: $N^{(\mathrm{true})} = 30$, $N^{(\mathrm{noise})} = 100$, $\alpha_0 = 0.00001$, $\alpha_1 = 0.0009$, $(\eta, \sigma) = \{(1, 0.1), (5, 1), (10, 2), (20,4), (50,6)\}$. For each pair $(\eta, \sigma)$, the simulations are repeated 1000 times and, in each repetition, the median relative mass error is calculated.

#### S.6 Validation of absolute relative mass error

A set of test masses $\mathbf{M}^{(\mathrm{test})} = \{m_1, m_2, \dots,\}$ are searched in the original dataset. The search is performed in the peaks list of each ROI pixel, individually, within a window of 20 ppm and 100 ppm for Orbitrap and TOF datasets, respectively. The set of test masses $\mathbf{M}'^{(\mathrm{test})} \subseteq \mathbf{M}^{(\mathrm{test})}$ found in at least 75% of the ROI size are considered for further analysis. We denote with the symbol $\mathbf{P}_m$ the set of pixels where the test mass $m$ is found.

We apply a further filter on the test masses similar to that employed for the pixel-wise recalibration regression models (Methods section). For each mass $m \in \mathbf{M}'^{(\mathrm{test})}$, we calculate the relative mass error in all pixels $p \in \mathbf{P}_m$ in ppm units (Eq. S6).

A Gaussian FFT KDE with Silverman bandwidth is used to estimate the density of the observed errors (expressed in ppm) of the masses in $\mathbf{M}'^{(\mathrm{test})}$. Therefore, we select the masses $\mathbf{M}''^{(\mathrm{test})} \subseteq \mathbf{M}'^{(\mathrm{test})}$ that have an error in ppm within the highest peak of the KDE density +/- 1 ppm. It is important to underline that no information about the range of the relative errors of the recalibration masses is used.

Once the filtered test mass set is generated, we calculate the absolute relative mass error $|\delta_{k,p}|$ (Eq. S6) from the original and recalibrated datasets. To have a one-to-one comparison, we consider the same match peaks for the original and recalibrated datasets. Finally, we calculate the median difference between the mass errors for each test mass in $\mathbf{M}''^{(\mathrm{test})}$ as the median of the difference between the absolute errors in the original and recalibrated peaks, $\Delta_k = \mathrm{median}\left(\boldsymbol{\gamma}_k^{(\mathrm{orig})} - \boldsymbol{\gamma}_k^{(\mathrm{recal})}\right)$, where $\boldsymbol{\gamma}_k = \left(|\delta_{k,p}|\right)$. We denote the vector of the median difference errors as $\boldsymbol{\Delta}' = (\Delta_k)$.

We, therefore, assess that the median of $\boldsymbol{\Delta}'$, denoted $\tilde{\Delta} = \mathrm{median}(\boldsymbol{\Delta}')$, is statistically different from zero by bootstrapping hypothesis testing.

The two-sided p-value is calculated as the proportion of times we observe a value more extreme than $\tilde{\Delta}$:

$$p = \frac{1}{B+1} \cdot \left(1 + \sum_{b=1}^{B} \mathbf{1}_{|\tilde{\Delta}_b| \geq |\tilde{\Delta}|}\right) \tag{S10}$$

where $\mathbf{1}$ represents the indicator function and $\tilde{\Delta}_b$ is the median of the bootstrapped values in replicate $b$. We set $B = 10000$.

Also, we estimate the 95% intervals of $\tilde{\Delta}$ using bootstrapping, corresponding to the 0.025 and 0.975 quantiles of the bootstrapped values.

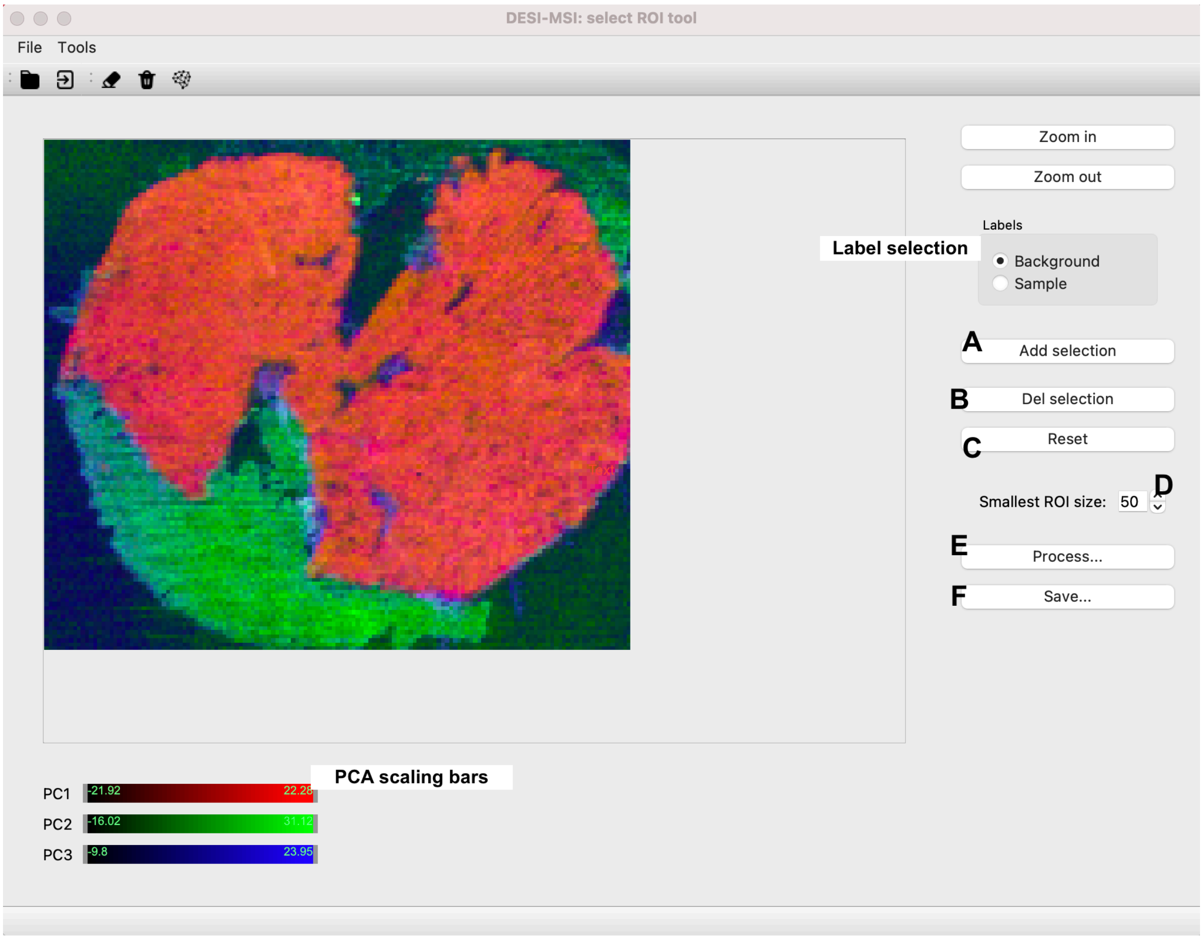


*Figure S 2 – A screenshot of the GUI for the selection of the sample ROI pixels. The scores of the first three principal components are plotted as RGB channels, adjusted in the slide bars below the image. The user can annotate (A, B, C) regions as "Background" and "Sample", after which a linear SVM model performs the segmentation of the entire image ('Process…',*

*E). The ROI is then saved in a CSV file in the same folder of the DESI-MSI data ('Save…', F). The connected regions smaller than the value in D are labelled as background.*

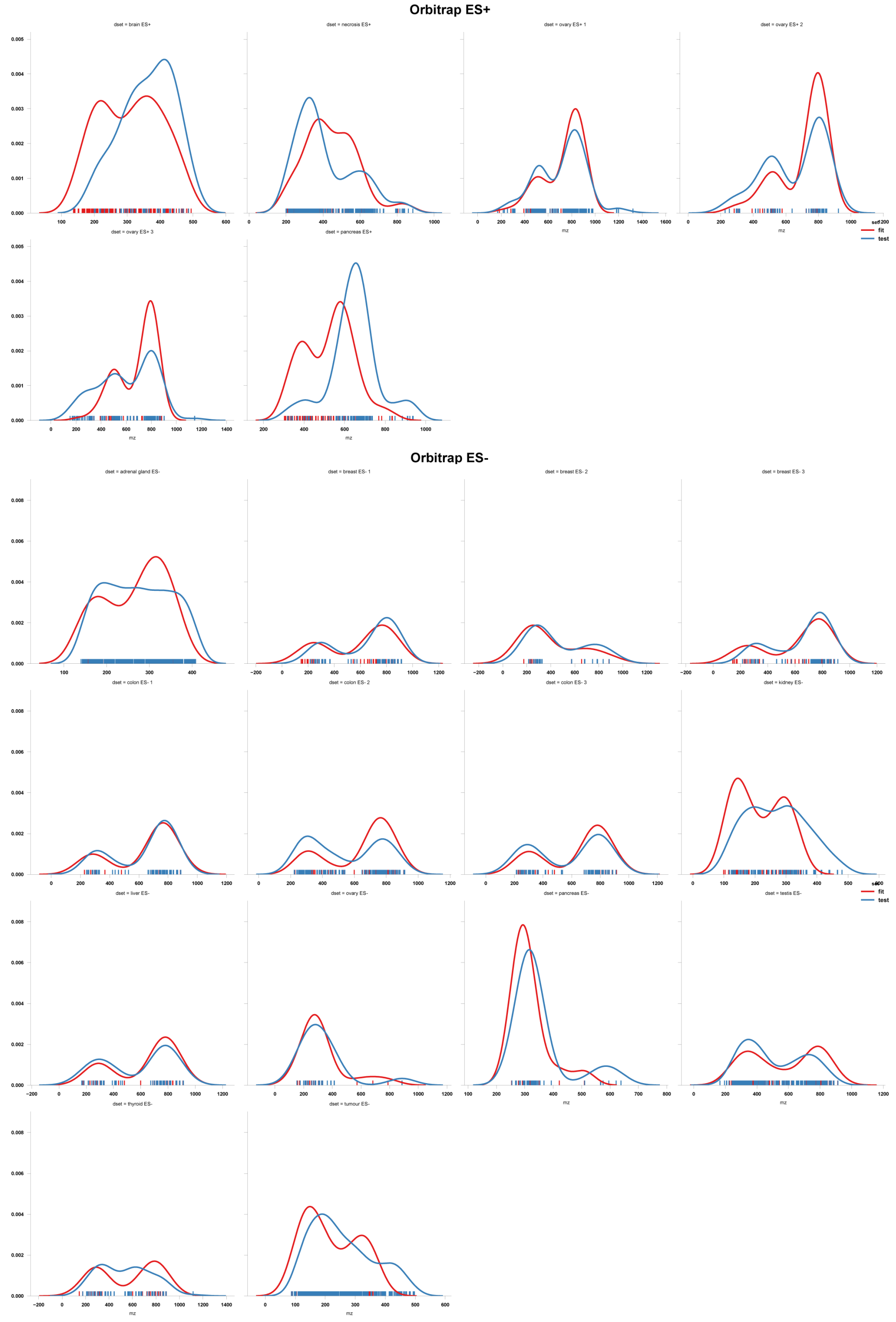


*Figure S 3 – Density plots of reference masses m/z values used for the pixel-wise regression models (red) and the external validation (blue) in the Orbitrap datasets. The plotted masses were used in at least 95% of the pixels of the dataset. Their distributions show that the reference masses covered the acquisition m/z range except for the pancreas ES- dataset, where*

*they span the values below 500 m/z. Also, the distributions of the test masses show that they covered the whole acquisition m/z range, thus providing a reasonable estimation of the errors after the recalibration. The densities were estimated using the function "distplot" available in the "seaborn" [10] package for Python.*

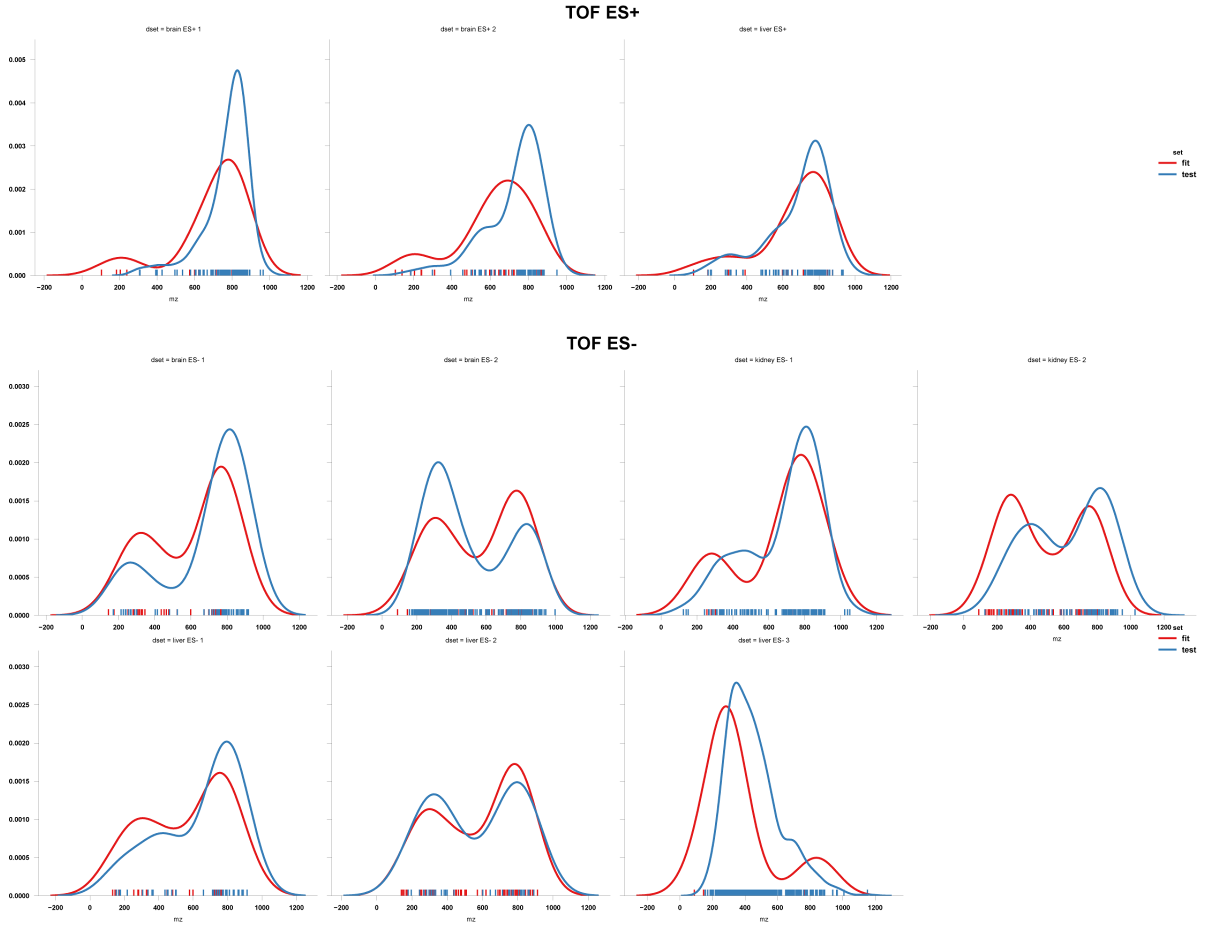


*Figure S 4 - Density plots of reference masses m/z values used for the pixel-wise regression models (red) and the external validation (blue) in the TOF datasets. The plotted masses were used in at least 95% of the pixels of the dataset. Their distributions show that the reference masses covered the acquisition m/z range except for the pancreas ES- dataset, where they span the values below 500 m/z. Also, the distributions of the test masses show that they covered the whole acquisition m/z range, thus providing a reasonable estimation of the errors after the recalibration. The densities were estimated using the function "distplot" available in the "seaborn" package for Python.*

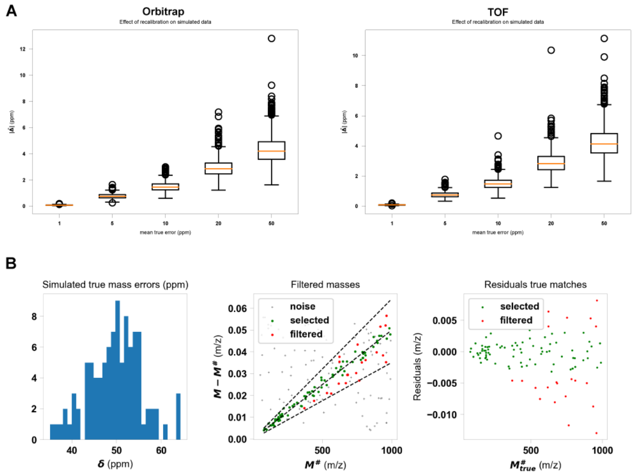


*Figure S 5 - Results of simulated pixel-wise recalibration models. Top: median error for the true matches using the Orbitrap model (left) and TOF model (right), with varying mean relative errors. Bottom: Example of simulated masses with a given distribution of relative errors (left). The masses' true matches are bounded as expected and scattered around the line corresponding to the mean of relative errors distribution. The selected points are plotted in green. The results confirm a reduction of the residual heteroskedasticity for the selected masses (green points, right). To increase the readability of the scatter plots, we simulated 100 true matches and 100 noise matches.*

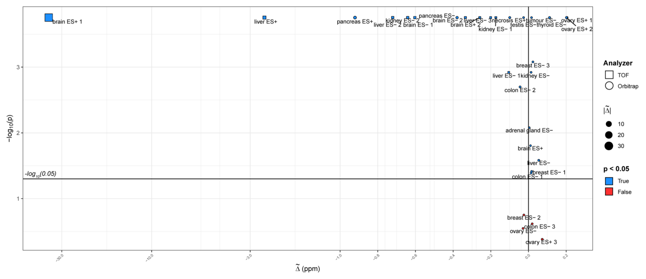


*Figure S 6 - Scatter plot representing the bootstrapping results for the median difference between the mass errors in the recalibrated datasets where the reference masses are modeled as a time series and those with independent models for each pixel. Each dataset is characterized by its $\tilde{\Delta}$ value (calculated following the method described in Section S6) and the significance value of the bootstrapping test. Negative values are interpreted as results favorable for the time series models.*

| **Dataset ORBITRAP** | $\tilde{\gamma}_0$ **(ppm)** | $\tilde{\gamma}_r$ **(ppm)** | $\tilde{\Delta}$ **(ppm)** | $(\tilde{\gamma}_0 - \tilde{\gamma}_r)^{(0.025)}$ **(ppm)** | $(\tilde{\gamma}_0 - \tilde{\gamma}_r)^{(0.975)}$ **(ppm)** | ***p-value*** |
|---|---|---|---|---|---|---|
| **adrenal gland ES-** | 0.248109 | 0.221925 | 0.043284 | 0.071297 | 0.180436 | **0.001059** |
| **brain ES+** | 0.34201 | 0.33341 | 0.004337 | -0.37296 | 0.289605 | 0.801828 |
| **breast ES- 1** | 0.420037 | 0.367504 | -0.00436 | -0.137 | 0.198306 | 0.9106 |
| **breast ES- 2** | 0.196908 | 0.389169 | -0.12603 | -0.31626 | -0.11299 | 0.327577 |
| **breast ES- 3** | 0.365379 | 0.297225 | 0.004484 | -0.04343 | 0.103513 | 0.581222 |
| **colon ES- 1** | 2.039641 | 0.245562 | 1.697414 | 1.597613 | 2.10267 | **0.000188** |
| **colon ES- 2** | 3.386558 | 0.383641 | 2.805863 | 2.74052 | 3.081262 | **0.000188** |
| **colon ES- 3** | 0.316833 | 0.24451 | 0.004068 | 0.007128 | 0.154435 | 0.0795 |
| **kidney ES-** | 2.28862 | 0.230932 | 1.993403 | 2.105863 | 2.292381 | **0.000188** |
| **liver ES-** | 1.001859 | 0.283977 | 0.488051 | 0.493124 | 0.689855 | **0.009474** |
| **necrosis ES+** | 1.044707 | 0.222903 | 0.719138 | 0.77699 | 0.961267 | **0.000188** |
| **ovary ES-** | 0.155214 | 0.166561 | 0.008538 | -0.06551 | 0.025438 | 0.060571 |
| **ovary ES+ 1** | 0.81531 | 0.772039 | 0.020406 | 0.069819 | 0.105741 | **0.000188** |
| **ovary ES+ 2** | 3.244488 | 1.247777 | 1.907598 | 1.793529 | 2.166747 | **0.000188** |
| **ovary ES+ 3** | 2.340457 | 1.09136 | 1.078912 | 1.003458 | 1.437133 | **0.000188** |
| **pancreas ES-** | 0.326273 | 0.274009 | 0.028505 | -0.02957 | 0.123313 | 0.19225 |
| **pancreas ES+** | 3.302328 | 2.635864 | 0.646432 | 0.657489 | 0.673178 | **0.000188** |
| **testis ES-** | 0.619615 | 0.52891 | 0.019138 | 0.007346 | 0.086667 | **0.000188** |
| **thyroid ES-** | 0.472414 | 0.235135 | 0.114365 | -0.04938 | 0.090732 | 0.706179 |
| **tumour ES-** | 1.410527 | 0.270817 | 0.996485 | 1.265094 | 1.341008 | **0.000188** |

*Table S 2 – Results of the comparison between the absolute test mass errors for the Orbitrap datasets. The first two columns represent the median value of the median error of the test masses in the original and recalibrated datasets, respectively. The third column represents the median of the difference between the median absolute errors in the original and recalibrated datasets, with their 0.025 and 0.975 quantiles estimated by bootstrapping. Finally, the last column contains the Benjamini-Hochberg corrected p-values obtained from the permutation test. The shading reflects the median increase (red) or decrease (green) of* $\tilde{\Delta}$.

| Dataset TOF | $\tilde{\gamma}_0$ (ppm) | $\tilde{\gamma}_r$ (ppm) | $\tilde{\Delta}$ (ppm) | $(\tilde{\gamma}_0 - \tilde{\gamma}_r)^{(0.025)}$ (ppm) | $(\tilde{\gamma}_0 - \tilde{\gamma}_r)^{(0.975)}$ (ppm) | ***p-value*** |
|---|---|---|---|---|---|---|
| **brain ES- 1** | 66.67604 | 2.959335 | 63.562 | 63.3035 | 64.27191 | **0.000188** |
| **brain ES- 2** | 0.972369 | 0.539609 | 0.114986 | 0.390499 | 0.614172 | **0.0025** |
| **brain ES+ 1** | 1.048839 | 0.633913 | 0.059468 | 0.214915 | 0.322758 | **0.03615** |
| **brain ES+ 2** | 66.97601 | 4.904221 | 62.02422 | 61.50941 | 62.56029 | **0.000188** |
| **kidney ES- 1** | 0.992241 | 0.937541 | -0.19181 | -0.12285 | 0.274135 | 0.178043 |
| **kidney ES- 2** | 1.742732 | 0.845695 | 0.296777 | 0.719461 | 0.781107 | **0.000188** |
| **liver ES- 1** | 1.22068 | 0.782113 | 0.093154 | -0.46368 | 0.877932 | 0.30408 |
| **liver ES- 2** | 44.53916 | 3.431842 | 36.46954 | 39.48972 | 41.86552 | **0.000188** |
| **liver ES- 3** | 2.348134 | 4.001103 | -1.31605 | -1.62528 | -1.56447 | **0.000188** |
| **liver ES+** | 15.23754 | 1.595811 | 12.78082 | 13.20028 | 13.91077 | **0.000188** |

*Table S 3 - Results of the comparison between the absolute test mass errors for the TOF datasets. The first two columns represent the median value of the median error of the test masses in the original and recalibrated datasets, respectively. The third column represents the median of the difference between the median absolute errors in the original and recalibrated datasets, with their 0.025 and 0.975 quantiles estimated by bootstrapping. Finally, the last column contains the Benjamini-Hochberg corrected p-values obtained from the permutation test. The shading reflects the median increase (red) or decrease (green) of* $\tilde{\Delta}$.
